# Comparing flickering and pulsed chromatic pupil light responses

**DOI:** 10.1101/2022.03.11.483966

**Authors:** Constanza Tripolone, Luis Issolio, Carlos Agüero, Alejandro Lavaque, Dingcai Cao, Pablo Barrionuevo

**Author notes:** Corresponding Author: Barrionuevo, P.

## Abstract

Protocols for chromatic pupil light reflex (PLR) testing considered mostly pulsed stimulation (pPLR). A more sophisticated and promising technique is based on the PLR to flickering stimulation (fPLR). Our aim was to compare fPLR and pPLR parameters in order to validate fPLR paradigm. Two different experiments were carried out in young participants to compare parameters of chromatic pupillary measurements under flickering and pulsed conditions. We found that the fPLR amplitude parameter was significantly associated with pPLR transient constriction parameter. Also, for some conditions, typical pulse parameters can be identified directly in the fPLR recordings.

## Introduction

The pupil light reflex (PLR) refers to the action of the iris muscles for modulating the pupil size as response to varying optical radiation. Classical way for assessing the PLR uses pulsed stimuli followed by a steady state. This type of method can be called as pulsed PLR (i.e., pPLR). The pPLR contains two clear behaviors: a transient or phasic response and a steady or tonic response. An alternative approach assessing PLR uses periodic flickering stimuli and therefore this type of method can be called as flickering PLR (fPLR) [1], and it is thought to reflect the transient state of the pupillary response (Fig. 1A). [2–5]

**Figure 1.**
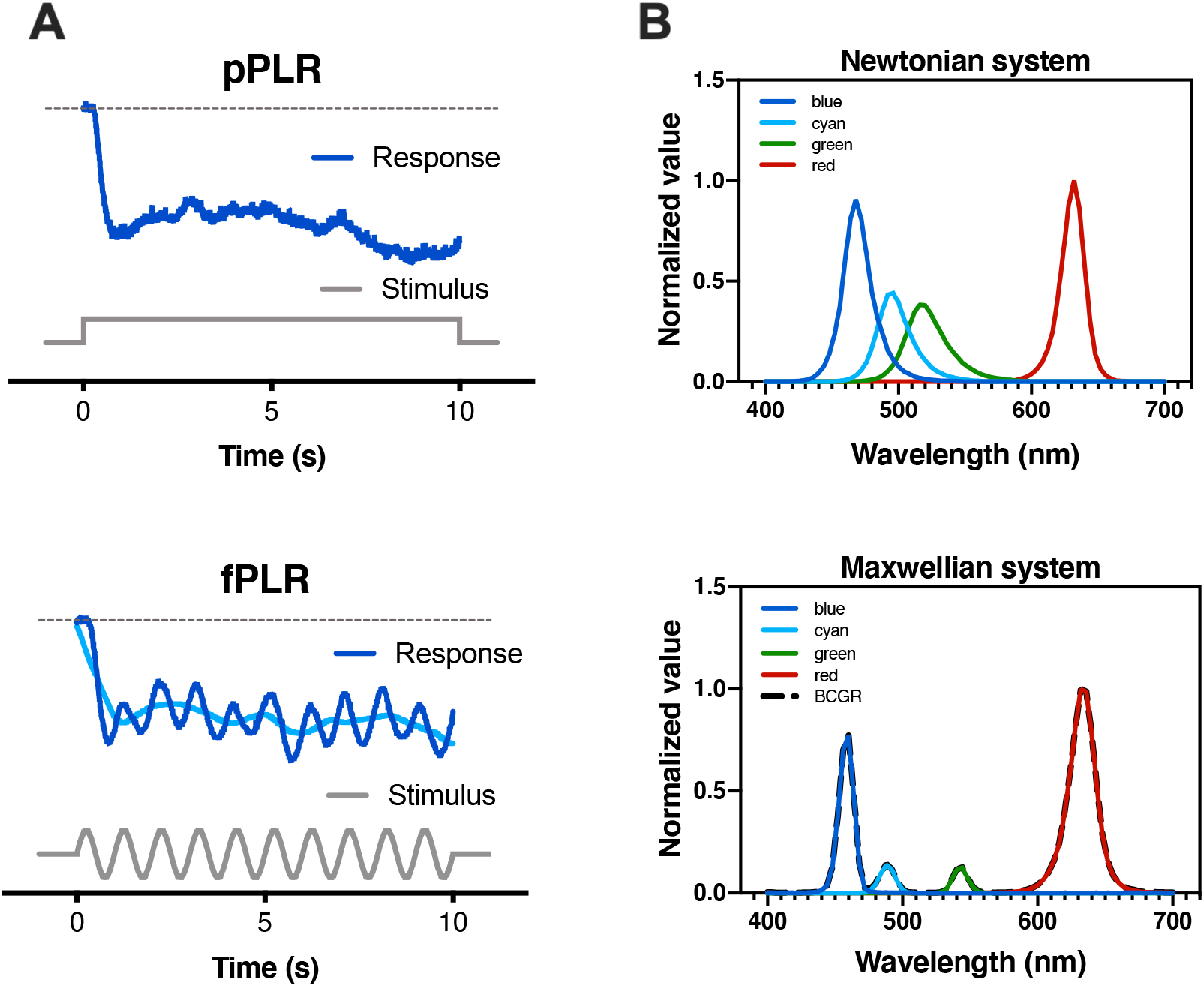
A) Pupillary responses to a pulsed stimulus, called pPLR (upper panel); and to a flickering stimulus, called fPLR, with the moving average data in light trace (lower panel). B) Spectra of the two apparatus configurations used in this study. The Newtonian system was used in Experiment 1 and the Maxwellian system was used in Experiment 2.

The PLR has been extensively used to study the underlying mechanisms in the pupillary control and to diagnose retinal diseases [6,7] However, in the last decades, the PLR study has gained renewed interest because the discovery of a small group of cells in the inner retina with photoreception capabilities, these cells are named intrinsically photosensitive retinal ganglion cells (ipRGCs). In mammals, the pupillary light reflex is almost entirely commanded by ipRGCs. [8] These cells were firstly reported in mice [9] and later in primates. [10] They express the photopigment melanopsin (peak sensitivity at ∼480nm) for photoreception [11], and receive inputs from outer retinal photoreceptors: rods and cones [10,12,13].

Another reason for interest in PLR is that it can serve as biomarker of the photoreceptors function. [14] Using monochromatic lights of different wavelengths, Gamlin and colleagues showed that melanopsin is the main contributor to pupilloconstriction for the post-illumination pupil response (PIPR) in humans. [15] Also, they showed that steady state of the pPLR in the dark-adapted eye is mainly maintained by action of melanopsin and rods. [16] While transient state is, depending on the light level and wavelength, mainly controlled by cones or rods. [2,17–19]

In recent years, different clinical studies have assessed retinal function using the PLR to monochromatic (usually blue and red) lights for enhancing the response to one photoreceptor above another. The non-invasive technique for assessing retinal behavior through measurements of pupil responses to different colored light stimulus was called chromatic pupillometry. [20,21] This technique is simple and fast and therefore is increasingly being used in investigative and clinical practice. It was employed with promising results in patients with, for example, retinitis pigmentosa, [17,22] glaucoma, [23–30] diabetes, [31,32] Leber congenital amaurosis, [17,33] and age-related macular degeneration. [34] It has produced the emergence of recommendations and protocols for the proper use of the pPLR technique. [14,17,21]

Most basic and clinical studies testing the PLR were carried out with pulsed stimulation in most cases. [7,14,21,35] On the other hand, the fPLR in combination with silent-substitution, a technique that allows each type of photoreceptor to be stimulated in isolation while maintains a constant adaptation in the other types of photoreceptor, [36–39] was used to determine that melanopsin, rods, L- and M-cones provide excitatory input to the pupil pathway, whereas S-cones provide inhibitory inputs. [3,19,38] In addition, recent studies showed an inhibitory participation for M-cones in the PLR. [40,41] Also, it has been shown that the fPLR to chromatic stimulus can serve to assess the interactions between outer and inner retinal photoreceptors. [1] One of the advantages of the fPLR is that noise inputs can be minimized through Fourier transformation for treatment in the frequency domain, especially if stimulation is sinusoidal. [2,3,5] Furthermore, fPLR and pPLR produce similar PIPRs, [1] therefore fPLR can serve to assess functioning of inner retinal cells. [15,42] However, the relationship between parameters of fPLR and parameters of pPLR has not been thoroughly studied.

The objective of this study was to compare fPLR and pPLR parameters in order to validate fPLR paradigm from well-known pPLR paradigm. Two different experiments were carried out to compare parameters of pupillary measurements under flickering and pulsed conditions.

## Materials & Methods

The two experiments were carried out with different experimental set-ups, stimulus size and procedures. Methods specificity are described in each Experiment Section.

### General Methods

#### Observers

8 young female observers (age: 35 ± 6 y.o.) and 8 young male observers (age: 35 ± 5 y.o.) were part of this study. All the observers had normal vision and were assessed with a visual acuity chart, and an optical coherence tomography system (RTVUE XR100-2, Optovue Inc., Fremont, CA), in order to ensure health of the retinal tissue; Decimal Visual Acuity: 1.0 ± 0.1, Retinal Nerve Fiber Layer Thickness (μm): 97.1 ± 10.1, Ganglion Cell Complex (μm): 97.4 ± 5.7. The study protocols were approved by the ethics committee of the Universidad Nacional de Tucumán, and were in accordance with the Declaration of Helsinki.

#### Stimuli conditions

For both instruments, the fPLR sinusoidal stimuli were generated with frequency of 1Hz. For the pPLR, the stimuli were “rectangular” pulses. The color conditions tested were: “Red”, in which only the red LED was turned on; “Blue”, in which only the blue LED was turned on; “Cyan”, in which only the cyan LED was turned on; “Green”, in which only the green LED was turned on; and “BCGR”, in which the blue, cyan, green, and red LEDs were turned on simultaneously (LEDs specificities are provided at table 1).

**Table 1.**
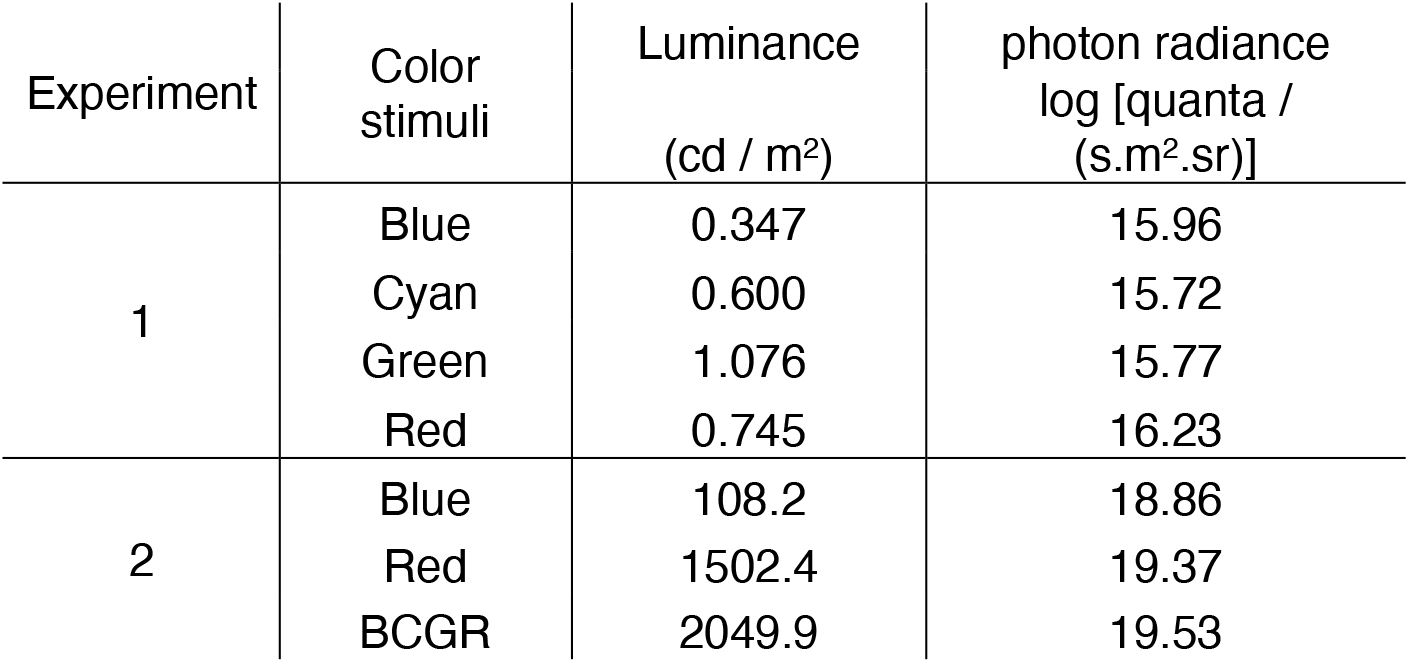
Luminance and photon radiance values at 50% of the maximum of each color stimuli condition for both experiments. Luminance in the Experiment 2 were computed from troland values divided the area of the artificial pupil in the Maxwellian view system.

| Experiment | Color stimuli | Luminance<br>(cd / m<sup>2</sup>) | photon radiance<br>log [quanta /<br>(s.m<sup>2</sup>.sr)] |
| --- | --- | --- | --- |
| 1 | Blue | 0.347 | 15.96 |
|  | Cyan | 0.600 | 15.72 |
|  | Green | 1.076 | 15.77 |
|  | Red | 0.745 | 16.23 |
| 2 | Blue | 108.2 | 18.86 |
|  | Red | 1502.4 | 19.37 |
|  | BCGR | 2049.9 | 19.53 |

#### Parameters

In the frequency domain, we proceeded to compute: 1) the amplitude (in millimeters) and 2) phase (in degrees) of the pupil response at 1Hz after Fourier transformation, following the procedure already reported for flickering responses [2]. In the temporal domain, we computed three parameters: 1) Time to minimum, which is the time from the start of the stimulation to achieve the first trough of the PLR, it is expresses in milliseconds; 2) Initial constriction, measured as the amplitude of the first trough of the constriction elicited after stimulus onset, it is expressed as percentage with respect to the baseline (defined as the averaged pupil diameter of the 100 frames previous the stimulus start); and 3) Plateau, which consisted in the average of the pupil diameter values from 6 s – 9 s after stimulus onset (Fig. 1A). Statistical analyses were carried out in STATA (StataCorp, USA).

### Experiment 1: Comparison between parameters of fPLR and pPLR

#### Rationale

It has been proposed that fPLR can give information about the transient component of pPLR. [2] Based on this assumption, parameters for the transient part of the pPLR (initial constriction and time to minimum) should be related to the parameters of the fPLR (amplitude and phase). In this experiment, we correlated parameters from fPLR with those from pPLR.

#### Apparatus

A Newtonian view arrangement used in this study was composed by a lab-made photostimulator-pupillometer system. The photostimulator consisted of four LEDs with different spectral composition as it is shown in the upper panel of Figure 1B and table 1. The LEDs intensities were controlled by an Arduino board following the premises introduced by a previous work. [38] The light of the four LEDs was carried out by a bundle of fiber optics to a monocular integrating sphere-like dome with a Lambertian inner surface. The observer’ eye was placed in front of the dome, in order to produce a Newtonian view. The camera was placed on the other end of the dome. We used a near-infrared camera (Manta G-031, Allied Vision Technologies GmbH, USA), which contains a CCD sensor with a temporal resolution range of (30 – 120) fps and spatial resolution of (656 × 492) pixels. Through a circular aperture the observer’s view fixed directly to the camera lens and surrounded by a uniform peripheral light field. A PC controlled both the Arduino board and the camera.

#### Calibration

Measurements of the LEDs spectrum were carried out every 4 nm with a spectroradiometer SpectraScan PR-715 (Photoresearch Inc, USA). Adapting background luminances (50% of maximum) were obtained with a luminancimeter L1009 (LMT GmbH, Germany), and belonged to the mesopic range (Table 1). Room light illuminance was obtained with a luximeter T-10M (Konica Minolta Inc, Japan)

#### Experimental conditions

We obtained pupillary recordings for Blue, Cyan, Green and Red stimuli conditions at 50% of amplitude (Exact light specification values are shown in Table 1) with respect to the adapting background. 15 observers participated at this experiment.

#### Procedure

Each observer was adapted during 5 minutes to the room light (10 lux) prior starting the recordings (since for this experiment the apparatus was located in a clinical setting; we could not darken completely the room). During this time, the system was adjusted in order to obtain the best image of the eye. For the pPLR paradigm, the observer looked into the dome to the camera and the recording started 10s prior to the stimuli in order to obtain the pupillary rest value. Then the pulsed stimulus was on during 5s. For fPLR, measurements were obtained with previous light adaptation to the 50% of the maximum of the specific illumination condition for 60 s. After that the sinusoidal waveform stimulus was presented during 10s. Recordings for each testing condition were repeated 3 times with an inter stimuli interval (ISI) of 30s. The observer was told to keep eyes open during the stimulus presentation and rest during the ISI. A session lasted less than 75 minutes per day to avoid fatigue. All sessions were carried out between 10AM and 6PM, avoiding testing up to one hour after lunch. Measurements in the fPLR paradigm were carried out in the same day as pPLR measurements but the order was randomized across observers and sessions.

#### Data Analysis

We obtained videos of the pupil during each trial at 90 fps for fPLR and 60 fps for pPLR. The video was fragmented in frames and an algorithm looked for circles at each frame in the position where the pupil was located. We obtained parameters of the PLR in postprocessing analysis in MATLAB (Mathworks Inc., USA) after averaging the repetitions. Repetitions were normalized recordings respect to the baseline for avoiding any long-term adaptation effect [43].

### Experiment 2: Prediction of pPLR parameters from fPLR parameters

#### Rationale

The pupil response in the fPLR paradigm usually appears mounted on a pupillary response that resemble the pPLR, as it is shown in the lower panel of Figure 1A (the light trace in the flickering response). We assessed how well the parameters of the pPLR agree with parameters obtained by the intrinsic pulsed response contained in fPLR recordings.

#### Apparatus

A Maxwellian view system was used in this experiment, it imaged the light of a photostimulator with four different LEDs (after band-pass filtering) at the pupil of the observer using a field achromatic lens and an artificial pupil with diameter of 2 mm. The spectral composition of the four LEDs is shown in the lower panel of Figure 1B and Table 1. The photostimulator control was a modification of the 5-primary system already reported in the literature. [38] The pupillary recording was consensual and carried out with an eye tracker (Cambridge Research System Ltd, Rochester, UK). Through the artificial pupil the observer saw a ring-shaped illuminated patch (outer border: 24°, inner border: 6°). The observer was instructed to fixate at the center of the ring.

#### Experimental conditions

We obtained responses for Blue, Red and BCGR stimuli conditions. For the pPLR, we tested four pulse amplitudes: 10%, 20%, 35%, and 50%, with respect to the adapting background. Same stimuli amplitudes were used for the fPLR. Three observers participated at this experiment.

#### Calibration

Measurements of the primaries’ spectrum were carried out every 4 nm with a spectroradiometer SpectraScan PR-715 (Photoresearch Inc, USA). Retinal illuminance was obtained following the method described elsewehere [44], from measurements with a luximeter pocket-Lux (LMT GmbH, Germany). Adapting background luminances were computed from retinal illuminance considering the artificial pupil and belonged to the photopic range (Table 1).

#### Procedure

The observer dark-adapted for 5 minutes. After that, a run consisted in light-adapting the observer to the 50% of the maximum of each illumination condition for 60 s or less, depending on the condition. Then, the stimulus was on during 10s and repeated 5 times with an ISI of 30s. During the ISI, the background was turned-on in steady 50% light adapting condition. The observer was told to keep eyes open during the stimulus presentation and rest during the ISI. A session lasted less than 75 minutes per day to avoid fatigue effects. All sessions were carried out between 10AM and 6PM, avoiding testing up to one hour after lunch.

#### Data analysis

the VideoEyetrace software provided by the manufacturer was used for obtaining pupil diameter at 250 fps. We also recorded the occurrences of an external trigger. This trigger was commanded by the computer that controls the LEDs intensities, and it was used for synchronization purposes. With the diameter of the pupil at each frame, we obtained parameters of the PLR in postprocessing analysis in MATLAB after averaging the repetitions.

To evaluate the steady-state response, we computed the Plateau parameter (see Parameters in General Methods Section). To assess the transient response, we computed time to minimum and initial constriction (%) parameters for both pPLR recordings and fPLR recordings (Fig. 1A). To compare parameters of pPLR and fPLR, we computed the difference in absolute terms of both PLR parameters (Eq. 1) and the “agreement variation” (Eq. 2).

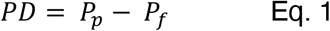

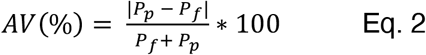

where, *PD* is the parameter difference, *P*_*f*_ is the parameter value of fPLR, *P*_*p*_ is the parameter value of pPLR and *AV* is an index of agreement variation.

## Results

### Experiment 1

Figure 2 shows the results for Experiment 1. Each row contains data for each colored LED stimuli condition (Blue, Cyan, Green and Red), for one stimulation amplitude (50%). Each data point corresponds to results for each participant. Initial pupil constriction from pPLR versus pupil amplitude for fPLR paradigm is shown in the left column. Time to minimum from pPLR versus fPLR phases are shown in the right column. Positive correlations were obtained for left column data for all stimuli condition. Instead, for right column data no correlation was found for most of the stimuli conditions. Simple linear regression Pearson ‘s correlation analyses showed that initial constriction parameter was significantly related to the amplitude parameter for Blue (*r* =0.77, *p* < 0.01), Cyan (*r* = 0.8, *p* < 0.001), Green (*r* = 0.88, *p* < 0.001), and Red (*r* = 0.87, *p* < 0.001). However, the time to minimum parameter could not be significantly associated with the phase parameter for Blue (*r* = 0.06, *p* = 0. 83), Cyan (*r* = -0.15, *p* = 0.62) and Green (*r* = -0.17, *p* =0.55). There was a weak correlation for Red (*r* = - 0.59, *p* = 0.03).

**Figure 2.**
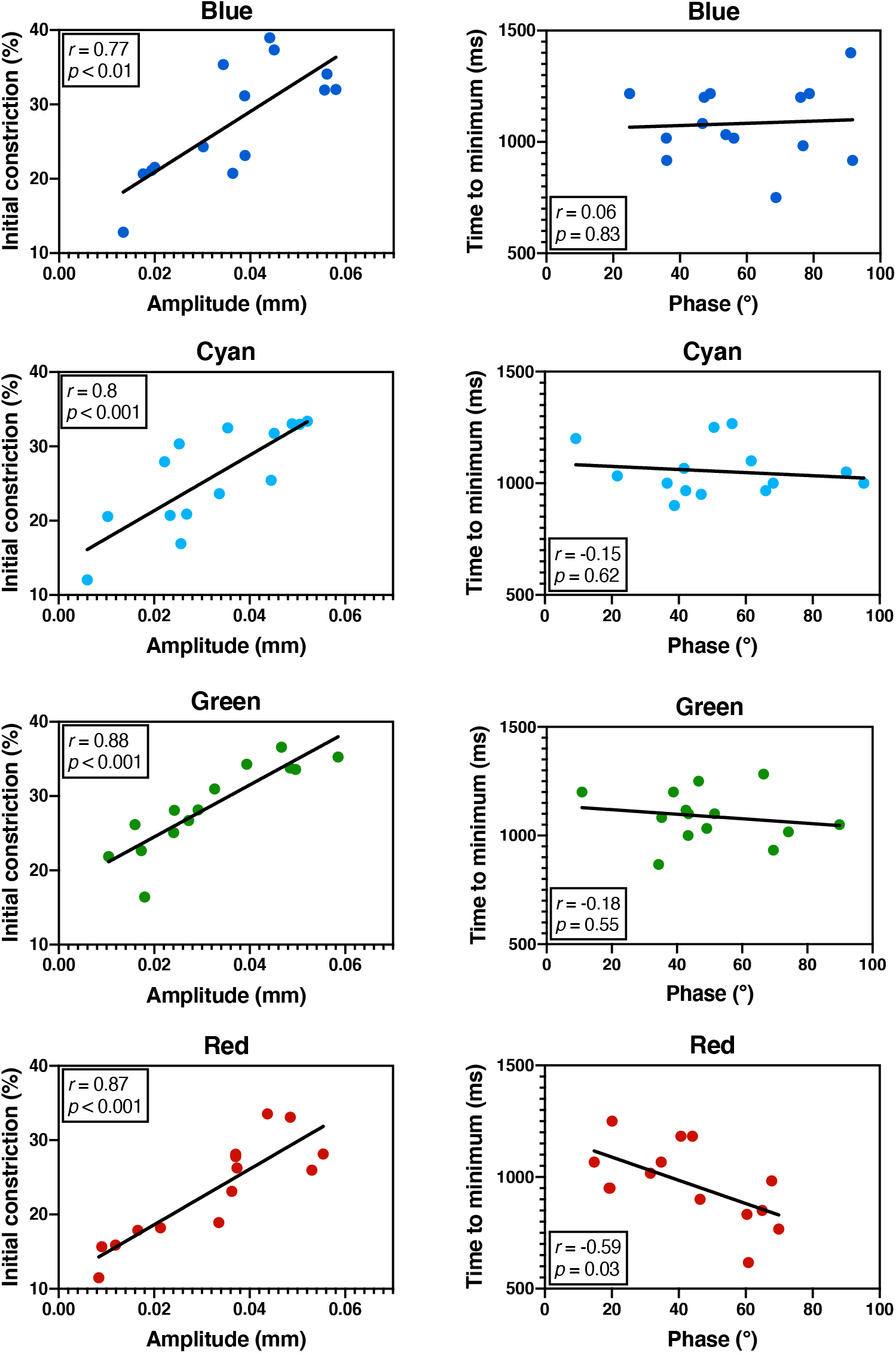
Results of Experiment 1 for each stimuli condition. Left column: Correlations between initial constriction of pPLR data and amplitude of fPLR data. Right column: Correlations between time to minimum of pPLR data and phase of fPLR data. Pearson’s correlation coefficient and statistical significance parameter are shown at each panel.

Thus, parameters considering the transient constriction intensity (Initial constriction for pPLR and Amplitude for fPLR) were related. Instead, temporal parameters (Time to minimum for pPLR and phase for fPLR) were not associated, except for Red stimuli.

### Experiment 2

Transient and steady-state parameters were obtained for three observers (Fig. S1). Comparison of pPLR parameters from pulsed recordings and flickering recordings is shown in Figure 3. Left column of Figure 3 contains the parameter differences for each observer and the color stimuli conditions as a function of the stimulation amplitude (Eq. 1). Since fPLR values were subtracted from pPLR values, a positive result indicates a higher value of the pPLR than fPLR, and a negative result indicate the opposite. Initial constriction values were mostly positives (66,6% for Blue, 83,3% for Red, and 75% for BCGR). Time to minimum values were also predominantly positives (100% for Blue, 75% for Red, and 91.6% for BCGR). Finally, at the bottom panels the plateau parameter is presented and the positive results are: 58.3% of the data for Blue stimuli, 58.3% of the data for Red stimuli, and 33.3% of the data for BCGR stimuli. ANOVAs considering as variable the type of paradigm (fPLR vs pPLR) showed significant differences for the Initial constriction (*df* = 1, *F* = 8.48, *p* < 0.01) and for the Time to minimum (*df* = 1, *F* = 18.84, *p* < 0.001), instead there was no difference for the Plateau (*df* = 1, *F* = 0.25, *p* = 0.62). Regarding the stimulation amplitude, there were significant differences for Initial constriction (*df* = 3, *F* = 90.78, *p* < 0.001) and for the Plateau (*df* = 3, *F* = 3.4, *p* < 0.05), however no significant differences were found for the Time to minimum parameter (*df* = 3, *F* = 1.69, *p* = 0.1846). Considering the color stimuli condition, we found significant differences for the Initial constriction (*df* = 2, *F* = 12.95, *p* < 0.001), and for the Plateau (*df* = 2, *F* = 51.52, *p* < 0.001). There was no significant difference for the Time to minimum parameter (*df* = 2, *F* = 3.22, *p* = 0.05).

**Figure 3.**
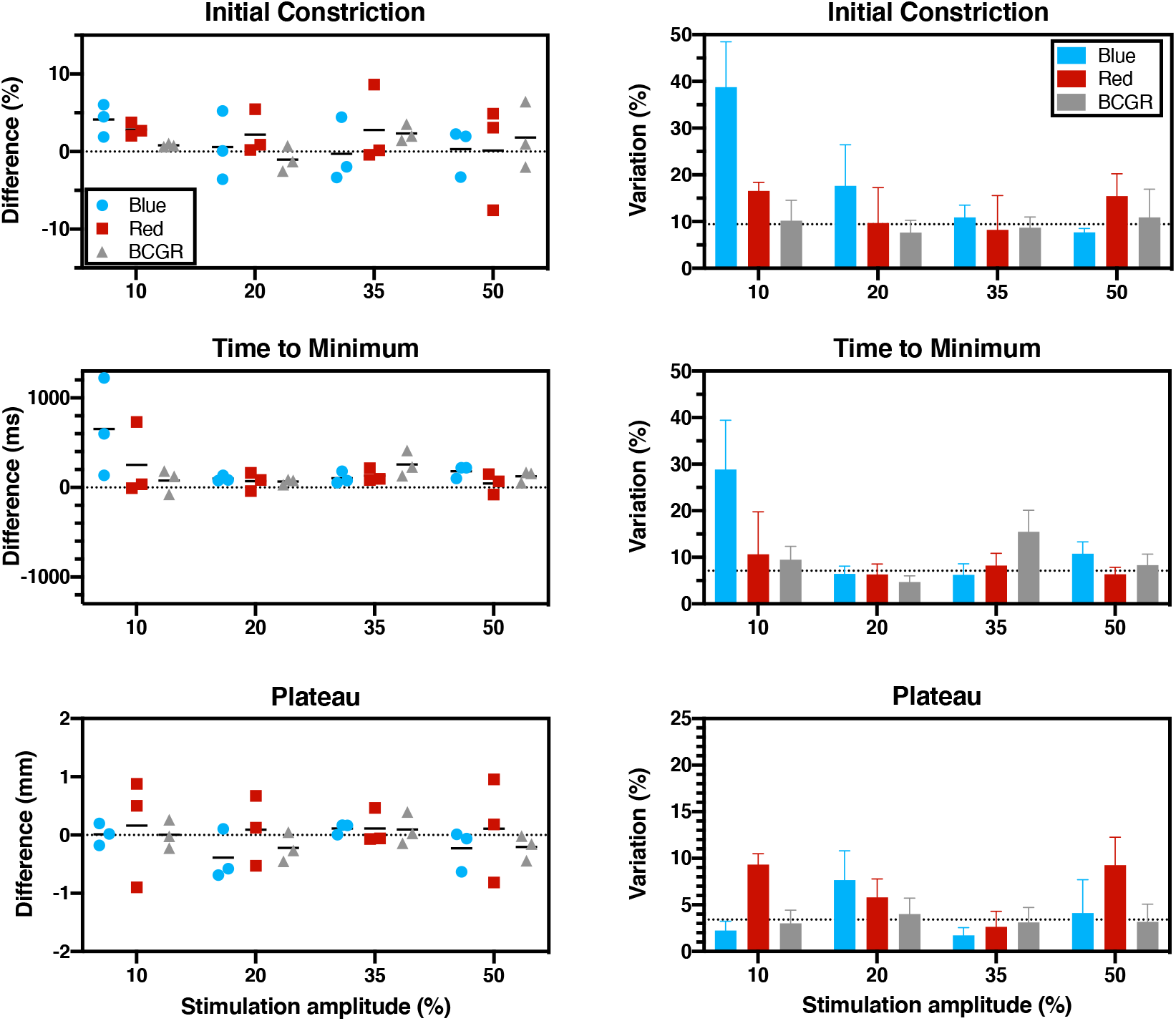
Results of Experiment 2. Differences for individual data (left column) and mean agreement variation (right column) between fPLR and pPLR parameters are shown for the four stimulation amplitudes and the three color stimuli. The horizontal black strokes in the left column represent the mean of observers ‘ data for each color stimuli at the respective stimulation amplitude. in Dashed lines in the right column represent the median of all data. Error bars are SEM.

The agreement variations between both sets of parameters are represented in right column of Figure 3. For the Initial constriction parameter, the agreement variation is lower for stimulation amplitude of 35%. In particular the difference is significant when compared with 10% (ANOVA: *df* = 3, *F* = 4.19, *p* < 0.05; Tukey post-test: *t* = -3.22, *p* < 0.05). Also, Blue data is significantly higher than BCGR values (ANOVA: *df* = 2, *F* = 4.02, *p* < 0.05; Tukey post-test: *t* = -2.78, *p* < 0.05). For the Time to minimum parameter, the agreement seems better for 20% and 50% of stimulation amplitude, although it was not confirmed with an ANOVA (*df* = 3, *F* = 2.87, *p* = 0.08). For the Plateau parameter, the agreement variation is small with respect to the other parameters; median = 3.4 (Plateau) vs 7.1 (Time to minimum) and 9.4 (Initial constriction). An ANOVA showed no significant differences for the color stimuli (*df* = 2, *F* = 3.22, *p* = 0.08), stimulation amplitude (*df* = 3, *F* = 1.63, *p* = 0.23) nor observer (*df* = 2, *F* = 0.58, *p* = 0.58).

For the steady-state response similar behavior for pPLR and fPLR was found according the agreement variation, particularly for a stimulation amplitude of 35% with respect to the baseline, in which small parameter differences were found all color stimuli. From transient parameters (Initial constriction and Time to minimum), it was shown that there is a higher and slower response of pPLR with respect to fPLR, which is evidenced in the high variations for these parameters (Fig. 3, right column).

## Discussion

We have assessed the relationship between pulsed and flickering pupil responses. We showed that the amplitude parameter obtained in the frequency domain of flickering responses was significantly associated with the transient constriction parameter obtained in the temporal domain of pulsed responses. However, the relationship of the phase parameter of flickering responses with the temporal parameter of the pulsed responses was weak or null. Also, we found that there were stimuli conditions where parameters of pulsed responses could be estimated analyzing same parameters for flickering responses. The correlations observed in the Experiment 1 between fPLR amplitude and pPLR Initial constriction suggest that fPLR is related to the transient component of the pPLR. The lack of correlation for temporal parameters avoids for confirming this conclusion could be related with the difference in adaptation between both tests (fPLR: preadaptation to 50% of each chromatic condition; pPLR: adaptation to room light), as a previous study also showed similar pattern between fPLR-derived latencies than pPLR latencies when measured for an adaptation light level range of at least 4 log units. [45,46]

From the nature of flickering responses, one could argue that steady-state parameters could not be computed using this paradigm. However, the analysis of the fPLR in the temporal domain showed that fPLR could also supply similar plateau responses than those of pPLR (Fig. 3 and Fig. S1). It means that besides the transient nature of flickering responses, this paradigm can give information of the PLR steady-state. Furthermore, previous studies showed that similar PIPR could be obtained from fPLR as from Pplr. [1,4] On the other hand, Gooley and colleagues showed a continuous increment in pupilar diameter (pupillary escape) for pPLR but not for fPLR. [47] This could cause discrepancies in the results of both paradigms in the sustained response (Plateau). We didn ‘t find that difference in our data possibly because the duration of our stimuli was only 10 s. Instead, for their results the difference between fPLR and pPLR started for stimuli durations higher than 24 s.

The minimum of pPLR was slower than the first constriction of fPLR (Experiment 2). This delay could be explained by the frequency of the fPLR (1Hz). It may happen that the frequency is not low enough for allowing the maximal possible initial constriction, since other pupillometry studies showed that maximal amplitude is achieved in lower frequencies [4,47–49]. In any case, for our data the time to minimum is independent of the stimulation amplitude in agreement with previous studies. [7,45] Also it is constant for color condition, stimuli type and observer (Fig. S1), Therefore, it is possible to compute this value to get an estimation of the time to minimum of the pPLR. Similar conclusion could be achieved for the amplitude of the constriction, therefore the initial constriction of pPLR could be estimated from fPLR amplitude (Fig. S1).

We showed that flickering responses can be analyzed in both temporal and frequency domains; adopting a classical electrical-engineering approach of two response components i.e., AC and DC components. [45,48,50] Therefore the DC component of the fPLR could be analyzed as a pPLR, and the AC component could provide additional information about the pupillary neural circuit. For example, it was previously showed that diffuse bipolar cells are responsible for double-frequency harmonic response to red-green modulation [51,52], and this result is evident for pupillary responses. [3] Our results contained this component for all color stimuli (Fig. S2). Therefore, we could speculate a role of diffuse bipolar cells in chromatic pupillometry however further testing is needed to corroborate this speculation.

The results of Experiment 1 and 2 were obtained in the mesopic and photopic region, respectively. Considering the weight of the photoreceptor contribution in fPLR is dependent of the light level as previous studies have shown, [1,2] our results for Experiment 1 would involve a higher contribution from rods than melanopsin-containing cells. Instead, the Experiment 2 would have higher melanopsin contribution with minimum rod contribution. This issue should not affect our conclusions since the approaches for study the fPLR are different in the two experiments. The analysis of the photoreceptor contributions across light levels was out of the scope of this study.

Considering the results of this study together with previous findings, we conclude that flickering responses can provide at least same information of neural processing as pulsed responses for healthy observers. Further comparisons in patients with ophthalmic and neural diseases could stablish the fPLR as a standard technique of chromatic pupillometry for use in clinical settings.

## Acknowledgments

CONICET PUE 0114 ILAV. Agencia I+D+i PICT 2019-03673. Constanza Tripolone acknowledges CONICET for the scholarship to carrying out her doctoral studies.

## Notes

### Competing Interest Statement

The authors have declared no competing interest.

