## Supplementary material for "Comparing flickering and pulsed chromatic pupil light responses": Supplemetary material

### Supplementary material of Manuscript: "Comparing flickering and pulsed chromatic pupil light responses".

Steady and transient parameter values of Experiment 2.

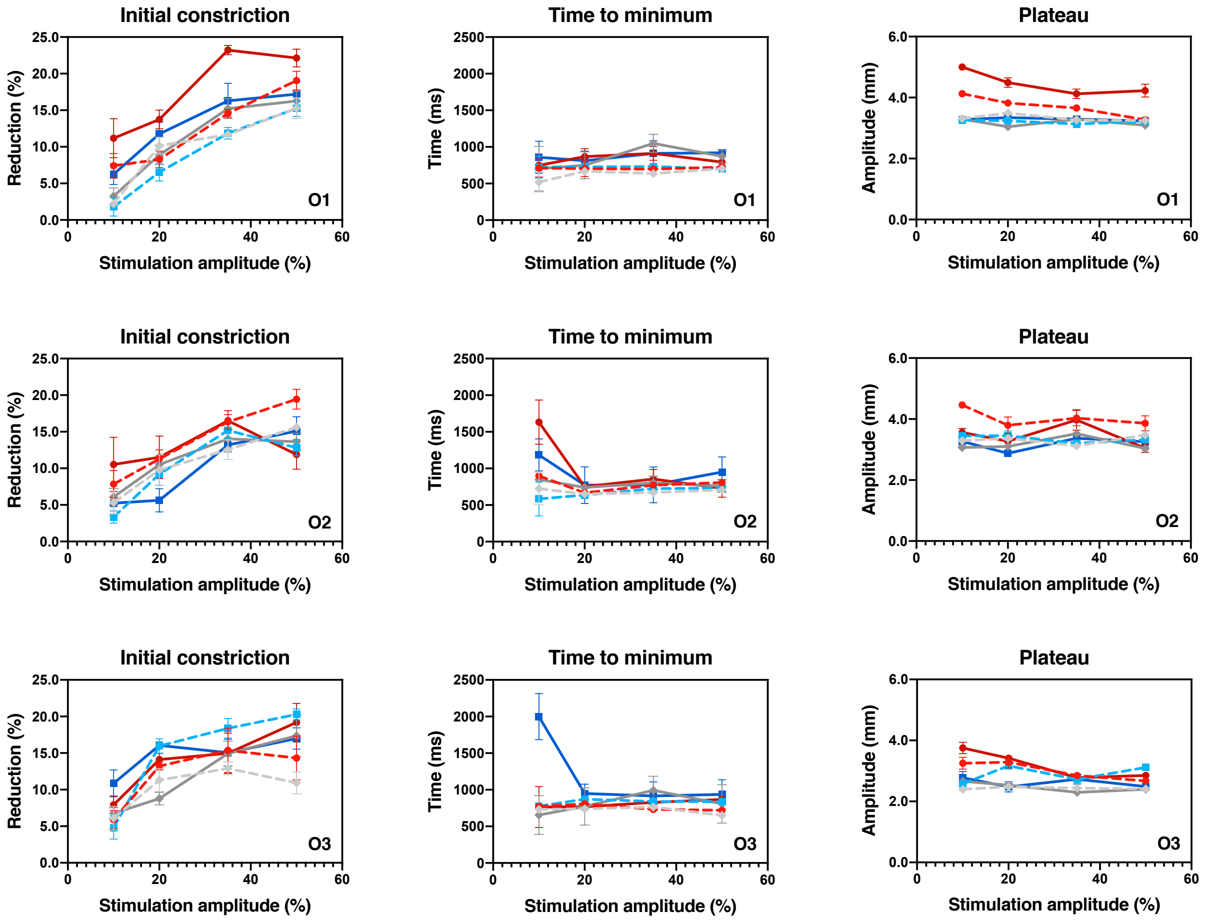


**Figure S1.** Parameter values for the transient component (Initial Constriction and Time to Minimum) and for the steady-state component (Plateau) of the pPLR (solid lines) and fPLR (dashed lines). Color stimuli data are shown in circle red dots (Red), square blue dots (Blue) and diamond gray dots (BCGR). Each row contains data for each observer. Error bars are SEM.

Relation of harmonics in Experiment 1

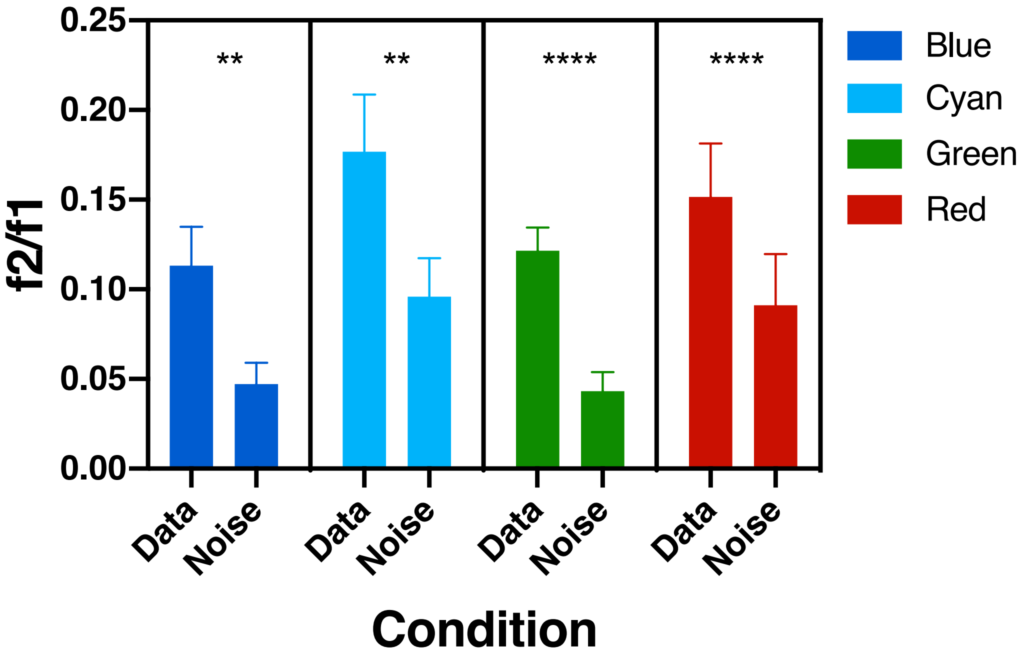


**Figure S2**. Relation of amplitude values (f2/f1) between component at 2Hz and component 1Hz (Data). The figure also contains the amplitude relation between the average of components around 2Hz (0.5 Hz – 0.8 hz & 1.2 Hz – 1.5 Hz) and the component at 1Hz (Noise). For all color stimuli the component in the second harmonics related with the first harmonic is higher and significative different than the pupil noise divided by the first harmonic. Error bars are SEM.
